# Genomic heterogeneity in pancreatic cancer organoids and its stability with culture

**DOI:** 10.1101/2022.07.03.498602

**Authors:** Olalekan Usman, Liting Zhang, Gengqiang Xie, Hemant M Kocher, Chang-il Hwang, Yue Julia Wang, Xian Fan Mallory, Jerome Irianto

## Abstract

The establishment of patient-derived pancreatic cancer organoid culture in recent years creates an exciting opportunity for researchers to perform a wide range of *in vitro* studies on a model that closely recapitulates the tumor. Among the outstanding questions in pancreatic cancer biology are the causes and consequences of genomic heterogeneity observed in the disease. However, to use pancreatic cancer organoids as a model to study genomic variations, we need to first understand the degree of genomic heterogeneity and its stability within organoids. Here, we used single-cell whole-genome sequencing to investigate the genomic heterogeneity of two independent pancreatic cancer organoids, as well as their genomic stability with extended culture. Clonal populations with similar copy number profiles were observed within the organoids, and the proportion of these clones was shifted with extended culture, suggesting the growth advantage of some clones. However, sub-clonal genomic heterogeneity was also observed within each clonal population, indicating the genomic instability of the pancreatic cancer cells themselves. Furthermore, our transcriptomic analysis also revealed a positive correlation between copy number alterations and gene expression regulation, suggesting the functionality of these copy number alterations.

## Introduction

Pancreatic ductal adenocarcinoma (PDAC) is the most common form of pancreatic cancer. With a 5-year survival rate of 11%, PDAC has one of the highest death rates among all cancers^1^. As part of the effort to improve the survival rate of PDAC and to develop more effective treatments, a better understanding of the disease biology is urgently needed. In recent years, multiple studies have reported success in establishing PDAC organoid culture from patient tumors^2–8^. These organoids were shown to closely recapitulate the phenotype, the genotype, and the drug response of their *in vivo* tumor counterparts. PDAC organoid culture provides researchers with the opportunity to perform various *in vitro* studies on a model that closely represents the *in vivo* tumor, which in turn, will allow for a broader, deeper, and more accurate understanding of the disease. One characteristic of PDAC is its genomic heterogeneity in both primary tumors and metastatic lesions^9–16^. However, the causes and consequences of the observed genomic heterogeneity are largely unknown. A better understanding of the genomic heterogeneity will help us elucidate both the etiology of the disease and its progression. PDAC organoid culture provides the opportunity to perform high-resolution genotyping and detailed mechanistic studies. Moreover, sample purity has been a significant issue in tissue-based genomic studies, where low purity has been shown to compromise the accuracy of the genomic data^17^. PDAC organoid culture, on the other hand, yields samples with the highest cancer cell purity.

To use PDAC organoids as a model to study genomic variations, however, we first need to understand the baseline genomic heterogeneity and stability of PDAC organoids. Without this understanding, it will be challenging to accurately interpret the genomic data after specific genetic modulations, treatments, or other perturbations. One of the pioneering studies by the Tuveson group demonstrated the feasibility and strategy of PDAC organoid culture using karyotyping to show the ploidy shifts within the extended organoid culture^3^. A following-up study by the same group derived single-cell clones from the organoid cultures^18^ and showed sub-clonal copy number variations (CNVs) between the clones, indicating genomic heterogeneity and instability within the PDAC organoids. Here, using single-cell whole-genome sequencing (scWGS) on two independent lines of PDAC organoids, we significantly increased the cellular resolution of organoid genotyping compared with previous studies. We observed genomic heterogeneity, in the form of chromosome copy number alterations, within both lines of PDAC organoids. Furthermore, we identified clonal and sub-clonal populations and elucidated their correlations through a phylogenetic tree analysis. Lastly, our transcriptomic analysis also revealed the functionality of these copy number alterations, suggesting a “gene dosage” effect^19–21^ of these copy number alterations that translates to gene expression regulation.

## Results

### Copy number alterations in two lines of patient-derived PDAC organoids

Two patient-derived primary tumor PDAC organoids (hPT1 and hPT2) were acquired and expanded for this study (**Figure 1A**). The doubling time of both organoids was about 7 days, hence the organoids were passaged at a 1:2 ratio weekly. Immunostaining clearly showed that the organoids were positive for both cytokeratin and epithelial cell adhesion molecule (EpCAM), reflecting their epithelial origin (**Figure 1B**). To analyze their chromosome copy number profiles, genomic DNA of both hPT1 and hPT2 organoids was subjected to single-nucleotide polymorphism microarray (SNPa), and an in-house algorithm was used to derive their copy number profiles at 1 Mb resolution (**Figure 1C**). We previously benchmarked this method and demonstrated that SNPa produced similar results as the comparative genome hybridization array^22^, which is the gold standard for chromosome copy number profile measurements^23^. Chromosome copy number alterations were observed across the genome of both hPT1 and hPT2 (Figure 1C). The total chromosome number was estimated from the summation of the average copy number of each chromosome. In general, copy gains were more prevalent than losses, resulting in a sub-triploid ploidy of 57.1 and 57 chromosomes for hPT1 and hPT2, respectively.

**Figure 1.**
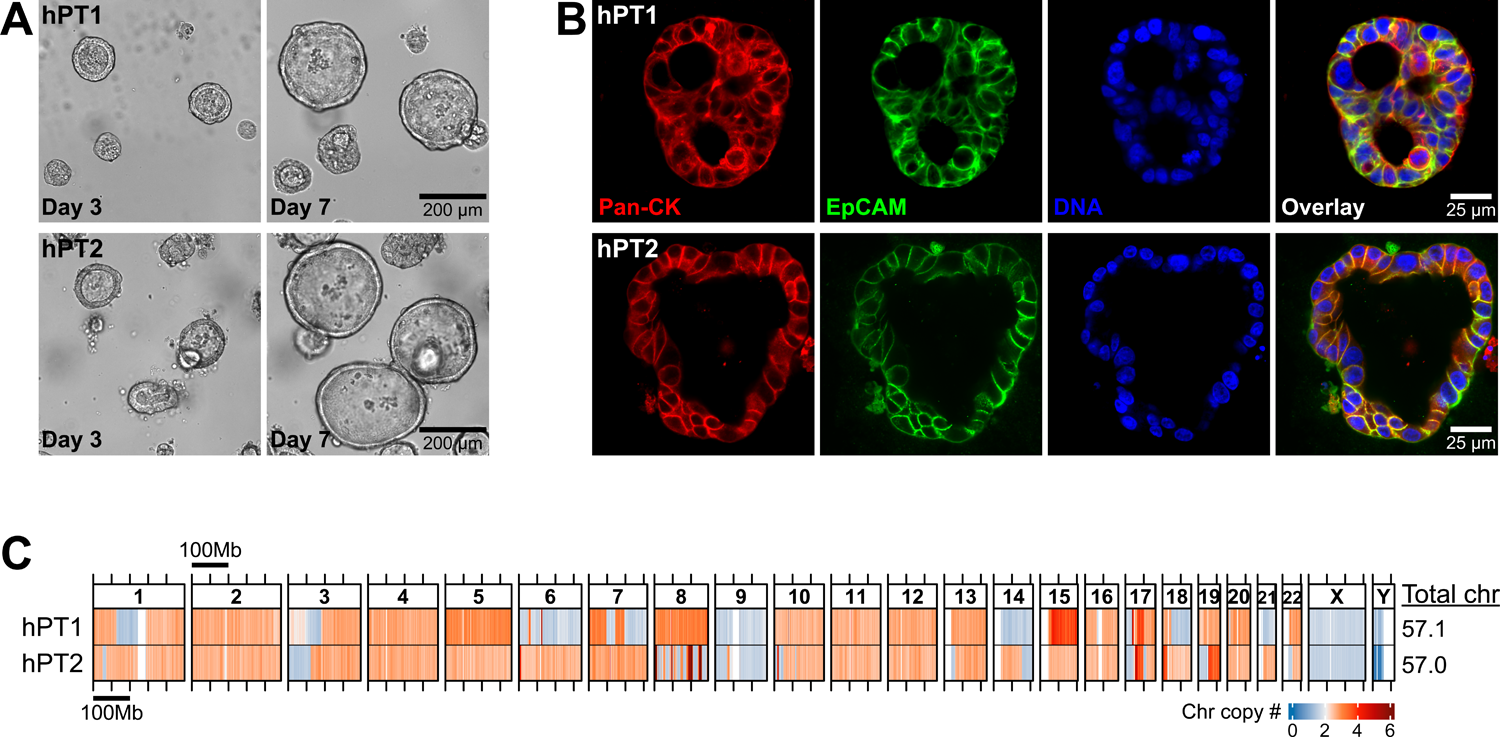
Chromosome copy number alterations in patient-derived pancreatic ductal adenocarcinoma (PDAC) organoids. (**A**) Representative images of primary tumor-derived PDAC organoids hPT1 and hPT2, from two independent patients, showing the growth of the organoids over 4 days (bar = 200 µm). (**B**) Confocal sections of hPT1 and hPT2 organoids immunostained for cytokeratin (red) and EpCAM (green), both are markers of epithelial and PDAC cells (bar = 25 µm). (**C**) Genomic DNA of hPT1 and hPT2 was subjected to SNPa and the resulting data was used to derive chromosome copy numbers of the organoids within 1Mb binning windows. The heatmaps illustrate chromosome copy numbers across the whole genome, where blue indicates chromosome loss (<2) and red indicates chromosome gain (>2). The total chromosome number was estimated from the summation of the average copy number of each chromosome. Chromosome copy number alterations were observed across the genome, and both organoids had sub-triploid ploidy of ∼57 chromosomes.

Additionally, we observed unusually high copy number alterations in chromosome 8 of hPT2, where the copy number appeared to oscillate between losses and very high gains. This pattern is in line with the typical signatures of chromothripsis, as reported in numerous cancers including PDAC^24, 25^. The role of chromothripsis in PDAC metastatic progression was suggested through the amplification of MYC at chromosome 8^26^.

### Genomic heterogeneity in PDAC organoids and genomic shift with prolonged culture

SNPa provides a bulk level overview of the average copy numbers of all the cells in the PDAC organoid. To improve our cellular resolution in understanding the genomic heterogeneity of PDAC organoids, we performed scWGS on both hPT1 and hPT2. We further examined each organoid line at two passage numbers to gain insights into the genomic stability of organoid culture.

From passage 3 of hPT1 organoids (hPT1 P3), we acquired scWGS data from 705 cells. We derived the copy number profile of each cell at 5Mb resolution. Of note, the resolution of the copy number profile from scWGS is lower than that from SNPa. Averaging the sequencing data at a bigger 5 Mb window is necessary due to the relatively shallow sequencing depth, ranging between 0.015X to 0.042X for the scWGS samples. To systematically analyze the genomic heterogeneity within each sample, we performed t-SNE analysis on the single-cell copy number profiles, followed by spectral clustering to derive clusters of closely related cells. The spectral clustering of the hPT1 P3 t-SNE plot resulted in five clusters (**Figure 2A**). Genomic heterogeneity was clearly observed within the hPT1 P3 population. Specifically, two distinct groups of cells were observed. The first group of cells was contained in Clusters 1 and 2, totaling 61 cells. Compared to the rest of the population, the cells in Clusters 1 and 2 had partial copy gains in chromosomes 1, 3, 7, 8, and 16 and whole chromosome copy gains in chromosomes 11, 12, 13, and 20. The second group of cells was contained in Clusters 3–5, totaling 644 cells. Genomic heterogeneity within this group was observed through the intermittent copy losses across the cells within chromosomes 1, 3, 6, 7, 9, 14, 18, 21, and X as well as copy gains within chromosomes 3, 8, 13, and 15. When we averaged the single-cell copy number data from the 705 hPT1 P3 cells, the resulting copy number profile is in accordance with the SNPa copy number profile of hPT1 (Figure 1C), confirming the concordance with the scWGS method. One difference between the aggregated scWGS profile and the SNPa profile is that the estimated number of chromosomes from the scWGS data is higher than the one estimated by SNPa. This slight discrepancy may be because they were derived from different methods and / or because of the different copy number resolutions, 5 Mb for scWGS and 1 Mb for SNPa. Nevertheless, the higher scWGS estimate is still within the sub-triploid ploidy range.

**Figure 2.**
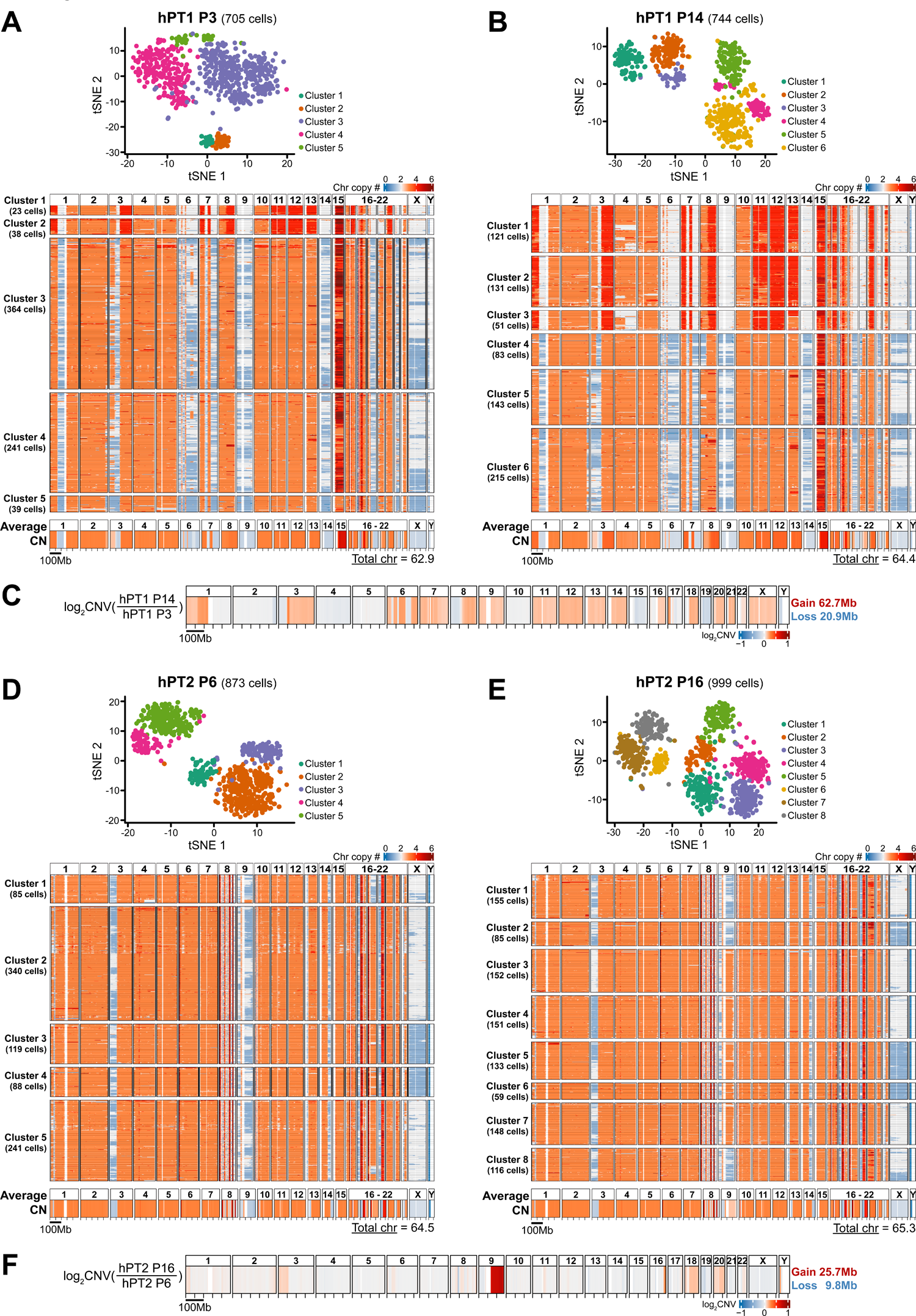
Single-cell whole-genome sequencing reveals genomic heterogeneity within PDAC organoids and the genomic shift with culture. (**A**) From passage 3 of hPT1 organoids (hPT1 P3), we acquired scWGS data of 705 cells. The single-cell copy number data derived from the scWGS were subjected to t-SNE analysis, followed by spectral clustering to derive the number of clusters within the sample. The spectral clustering of the hPT1 P3 t-SNE plot resulted in five distinctive clusters. The single-cell chromosome copy number profiles of each cluster were plotted in form of heatmaps, with the resolution of a 5 Mb binning window. Genomic heterogeneity is evident within the population. Clusters 1 and 2 show a distinct group of cells (totaling 61 cells) with partial chromosome copy gains in chromosomes 1, 3, 7, 8, and 16, in addition to whole chromosome copy gains in chromosomes 11, 12, 13, and 20. Genomic heterogeneity can also be observed in the rest of the cells (Clusters 3–5, totaling 644 cells) through the intermittent copy losses within chromosomes 1, 3, 6, 7, 9, 14, 18, 21, and X, as well as intermittent copy gains within chromosomes 3 and 15. The bottom heatmap shows the averaged copy number from the 705 hPT1 P3 cells. The copy number pattern across the genome is in accordance with the hPT1 copy number profile derived from the SNPa in Figure 1C. The total number of chromosomes was estimated to be 62.9, which is slightly higher than the one estimated by the SNPa, but both of them are still within the sub-triploid ploidy range. (**B**) From passage 14 of hPT1 organoids (hPT1 P14), we acquired scWGS data of 744 cells. The spectral clustering of the hPT1 P14 t-SNE plot resulted in six distinctive clusters. The copy number profiles of hPT1 P14 Clusters 1–3 resemble the cells in hPT1 P3 Clusters 1 and 2, however, the proportion of these cells increases within the hPT1 P14 population (totaling 303 cells), suggesting a clonal population expansion with the extended culture of 11 passages. Genomic heterogeneity within the hPT1 P14 Clusters 1–3 cells can be observed in chromosomes 1, 4, 5, 11, 13, 16, and 22. The copy number profiles of the rest of the cells (Clusters 4–6, totaling 441 cells) look similar to the hPT1 P3 Clusters 3–5 cells, except for the partial loss of chromosome 8 within all of the hPT1 P14 Cluster 4 cells and some cells in Clusters 5 and 6. The bottom heatmap shows the averaged copy number from the 744 hPT1 P14 cells. The total number of chromosomes was estimated to be 64.4, which is higher than hPT1 PT3, indicating copy number gains with extended culture. (**C**) The CNV between hPT1 P3 and hPT1 P14 was derived from the averaged copy number data. Chromosome copy gains were observed in chromosomes 1, 3, 6, 7, 8, 9, 11, 12, 13, 14, 16, 17, 18, 20, 21, and X, which correspond to the clonal expansion in hPT1 P14 Cluster 1–3. All combined, the extended culture resulted in approximately 62.7 Mb copy gains and 20.9 Mb copy losses. (**D**) From passage 6 of hPT2 organoids (hPT2 P6), we acquired scWGS data of 873 cells. The spectral clustering of the hPT2 P6 t-SNE plot resulted in five distinctive clusters. Similar to hPT1 organoids, genomic heterogeneity can be observed within the hPT2 population. In contrast to the other clusters, the cells within Cluster 1 have intermittent copy losses in chromosome 18, instead of copy gains. Eleven cells within Cluster 1 have uniquely partial gains in chromosome 9 and partial diploid in chromosome 4. The copy number profiles of the rest of the cells (Clusters 2–5, totaling 788 cells) are generally similar to each other, with some level of genomic heterogeneity in form of intermittent copy losses within chromosomes 3, 9, 13, 14, 17, 19, and X. The bottom heatmap shows the averaged copy number from the 873 hPT2 P6 cells, which is in accordance with the hPT2 copy number profile derived from the SNPa in Figure 1C. The total number of chromosomes was estimated to be 64.5. (**E**) From passage 16 of hPT2 organoids (hPT2 P16), we acquired scWGS data of 999 cells. The spectral clustering of the hPT2 P16 t-SNE plot resulted in eight distinctive clusters. The copy number profile of hPT2 P16 Clusters 1, 2, and 8 (totaling 356 cells) resemble the cells in hPT2 P6 Clusters 2–5, with the addition of copy gains in chromosome 20 in some cells. The rest of the cells in Clusters 3–7 (totaling 643 cells) have partial copy gains in chromosome 9, resembling the 11 cells in hPT2 P6 Cluster 1, again suggesting a clonal population expansion with the extended culture. However, in contrast to the cells in hPT2 P6, these 643 cells also have copy gains in chromosomes 4 and 18. The bottom heatmap shows the averaged copy number from the 999 hPT2 P16 cells. The total number of chromosomes was estimated to be 65.3, which is higher than hPT2 P6, indicating copy number gains with extended culture. (**F**) The CNV between hPT2 P6 and hPT2 P16 was derived from the averaged copy number data. Significant chromosome copy gains were observed within chromosome 9, which were caused by the clonal expansion in hPT2 P16 Clusters 3–7. Other copy gains were also observed in chromosomes 3, 16, 18, and 20. The extended culture resulted in approximately 25.7 Mb copy gains and 9.8 Mb copy losses.

Next, we performed scWGS analysis on passage 14 of hPT1 organoids (hPT1 P14). As we passaged the organoids weekly, hPT1 P3 was cultured for 11 weeks to reach hPT1 P14. At P14, we acquired scWGS data from 744 cells. The spectral clustering of the hPT1 P14 t-SNE plot resulted in six clusters (**Figure 2B**). From the copy number profiles, two distinct groups of cells were also observed in hPT1 P14. The copy number profiles of the first group of cells in Clusters 1–3 resembled the cells in hPT1 P3 Clusters 1 and 2. However, the proportion of these cells in hPT1 P14 grew to ∼40.7% (303 cells out of 744 cells) from ∼8.7% in hPT1 P3, suggesting a clonal population expansion with extended culture. Genomic heterogeneity within the hPT1 P14 Clusters 1–3 cells was observed within chromosomes 1, 4, 5, 11, 13, 16, and 22, which were absent in hPT1 P3 Clusters 1 and 2 cells. The copy number profiles of the 441 cells in hPT1 P14 Clusters 4–6 resembled the hPT1 P3 Clusters 3–5 cells, with the exception of the partial loss of chromosome 8 within all of the hPT1 P14 Cluster 4 cells and some of the cells in Clusters 5 and 6. In all, the total number of chromosomes was estimated to be 64.4, which is slightly higher than in the hPT1 P3, indicating copy number gains with the extended culture. Indeed, when we derived the CNV between hPT1 P3 and P14 from the averaged copy number profile (**Figure 2C**), 62.7 Mb copy gains were observed across chromosomes 1, 3, 6, 7, 8, 9, 11, 12, 13, 14, 16, 17, 18, 20, 21, and X and 20.9 Mb copy losses were observed in other chromosomes. Most of the observed genomic shifts corresponded to the clonal expansion in hPT1 P14 Clusters 1–3.

Similar scWGS analyses were performed for the hPT2 organoids. From passage 6 of hPT2 organoids (hPT2 P6), we acquired scWGS data from 873 cells. The spectral clustering of the hPT2 P6 t-SNE plot resulted in five clusters (**Figure 2D**). Three distinct groups of cells were observed within the single-cell copy number profiles of hPT2 P6 cells. The differences between these three groups of cells revolved around chromosomes 4, 9, and 18. First, the 85 cells within Cluster 1 had intermittent copy losses in chromosome 18, instead of the copy gains that were observed in the other clusters. Second, within Cluster 1, 11 cells had uniquely partial gains in chromosome 9, accompanied by partial diploid in chromosome 4. The rest of the 788 cells in Clusters 2–5 had copy gains in chromosomes 4 and 18, and intermittent losses in chromosome 9. Genomic heterogeneity in these 788 cells was observed in form of intermittent copy losses within chromosomes 3, 9, 13, 14, 17, 19, and X. The unusual oscillating copy number profile in chromosome 8, as illustrated in Figure 1C, was also observed under scWGS in all cells. Furthermore, the averaged copy number profile of hPT2 P6 under scWGS also highly resembled the SNPa copy number profile of hPT2 P6 in Figure 1C. The total number of chromosomes for the hPT2 P6 cells was estimated to be 64.5, again, slightly higher than the number estimated by the SNPa copy number profile (Figure 1C). After an extended culture for 10 additional passages, the scWGS data of hPT2 P16 revealed a clonal expansion of the cells that had partial copy gains in chromosome 9, overtaking the population majority from ∼1.3% in hPT2 P6 to ∼64.4% in hPT2 P16 (Clusters 3–7, totaling 643 cells out of 999 cells, **Figure 2E**), again suggesting a clonal population expansion with extended culture. However, in contrast to hPT2 P6, these 643 hPT2 P16 cells had copy gains in both chromosomes 4 and 18. The rest of the 356 cells in hPT2 P16 Clusters 1, 2, and 8 resembled the cells in hPT2 P6 Clusters 2–5, with the addition of copy gains in chromosome 20 in some cells. From the averaged copy number profile, we estimated the chromosome number of hPT2 P16 to be 65.3, which was slightly higher than the number estimated for hPT2 P6, again suggesting a gain of copy number with extended culture. Indeed, from the CNV analysis between hPT2 P6 and P16 (**Figure 2F**), significant copy gains were observed within chromosome 9, which were caused by the clonal expansion in hPT2 P16 Clusters 3–7, and also other copy gains in chromosomes 3, 16, 18, and 20. All combined, the extended culture resulted in approximately 25.7 Mb of copy gains and 9.8 Mb of copy losses.

Next, to investigate the evolutionary distance between the clusters of cells, the copy number profiles of each cluster were averaged and used to derive phylogenetic trees for hPT1 and hPT2. The maximum parsimony method^27^ was used to build the trees from the early passage clusters, followed by the minimal distance pairings of the late passage clusters. This approach enabled us to reveal the correlation between the clusters of cells from different culture periods and to quantify the copy number differences between the clusters, as a measure of genomic heterogeneity. The hPT1 phylogenetic tree revealed two distinct groups of clusters that were separated by ∼1500 Mb of copy number differences (**Figure 3A**). Closer to the diploid root, the first group includes hPT1 P3 Clusters 3-5 (P3 C3–5), and hPT1 P14 Clusters 4-6 (P14 C4-6). P14 C5-6 are paired to P3 C4, while P14 C4 is paired to P3 C5. These pairings indicate the correlation between the late passage clusters to the early ones, and again confirm the existence of the same clonal population across the extended culture. The estimated chromosome number ranges from 56.1 to 62.8. The other group of clusters includes P3 C1-2, and P14 C1–3, which are paired to P3 C1. The estimated chromosome number ranges from 67.3 to 71.1. Unlike the first group, the clusters in this second group spanned within ∼500 Mb of copy number differences, indicating a lower level of genomic heterogeneity within these clusters.

**Figure 3.**
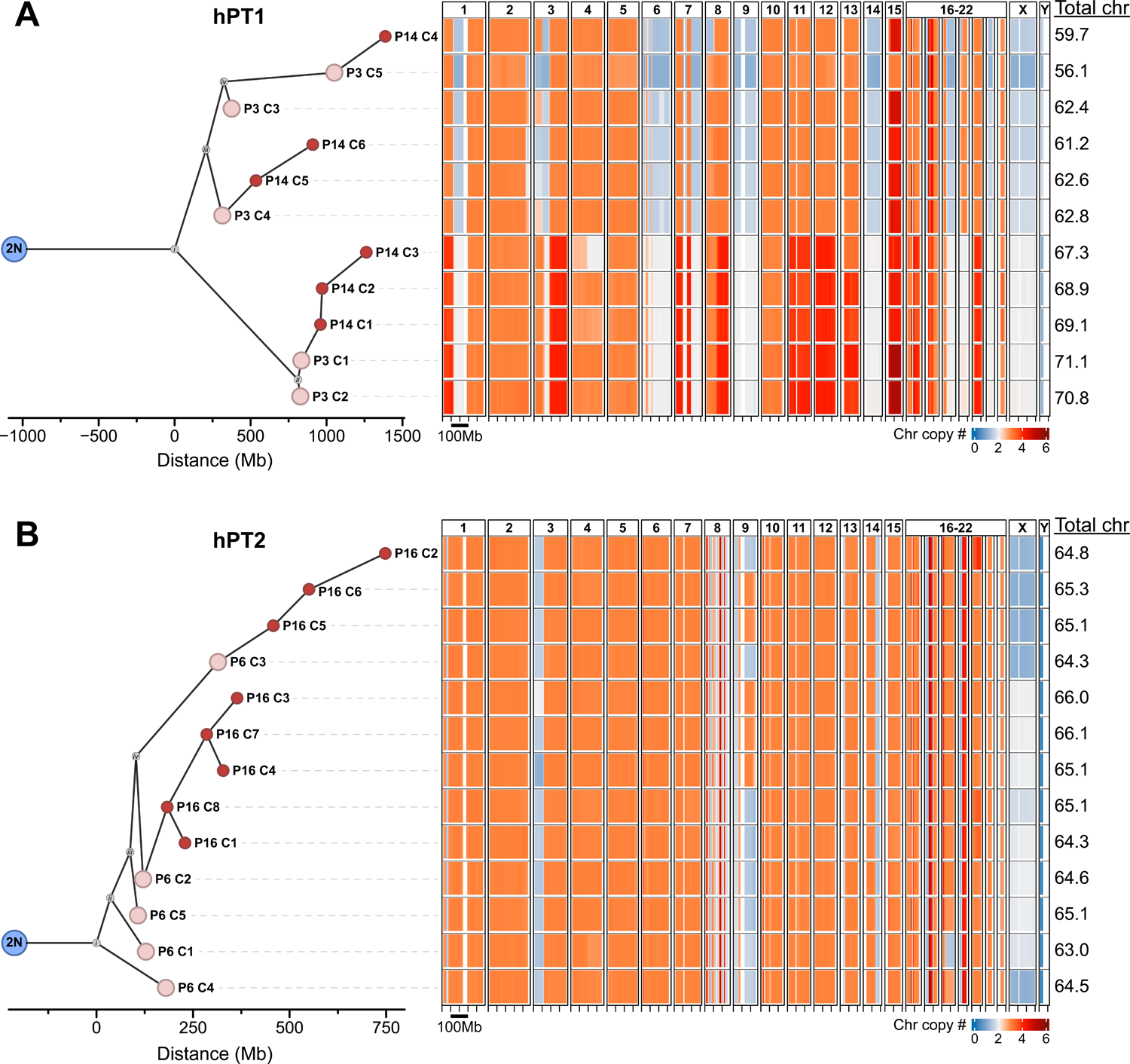
Phylogenetic tree analysis reveals the correlation between the identified sub-population clusters and enables the quantification of genomic heterogeneity. The single-cell chromosome copy number data of the clusters identified in Figure 2 were averaged and used to derive phylogenetic trees for hPT1 and hPT2. The maximum parsimony method was used to build the trees from the early passage clusters, followed by the minimal distance pairings of the late passage clusters. These trees reveal the correlation between the clusters of different culture periods and quantify the copy number differences between the clusters, as a measure of genomic heterogeneity. (**A**) hPT1 phylogenetic tree reveals two distinct groups of clusters that are separated by ∼1500 Mb of copy number differences. Closer to the diploid (2N) root, the first group includes hPT1 P3 Clusters 3-5 (P3 C3–5), and hPT1 P14 Clusters 4-6 (P14 C4-6). P14 C5-6 are paired to P3 C4, while P14 C4 is paired to P3 C5. The estimated chromosome number ranges from 56.1 to 62.8. The other group of clusters, spread within ∼500 Mb of copy number differences, includes P3 C1-2, and P14 C1–3, which are paired to P3 C1. The estimated chromosome number ranges from 67.3 to 71.1. (**B**) The clusters in the hPT2 phylogenetic tree are distributed within ∼750 Mb of copy number differences, which is smaller than the hPT1 tree, suggesting lower genomic heterogeneity within the hPT2 population. The hPT2 P6 Clusters 1-5 (P6 C1-5) are spread within ∼300 Mb and are closely related to each other. hPT2 P16 Clusters 1, 3, 4, 7, and 8 (P16 C1, C3-4, C7-8) are paired to P6 C2, while P16 C2 and C5-6 are paired to P6 C3. The estimated chromosome number ranges from 63 to 66.1.

The clusters in the hPT2 phylogenetic tree were distributed within ∼750 Mb of copy number differences, which was smaller than the hPT1 tree, suggesting lower genomic heterogeneity within the hPT2 population (**Figure 3B**). The hPT2 P6 Clusters 1-5 (P6 C1-5) are spread within ∼300 Mb and are closely related to each other. hPT2 P16 Clusters 1, 3, 4, 7, and 8 (P16 C1, C3-4, C7-8) are paired to P6 C2, while P16 C2 and C5-6 are paired to P6 C3. The estimated chromosome number ranges from 63 to 66.1. The copy number changes associated with each edge of the phylogenetic trees can be found in **Table 1**.

**Table 1.**
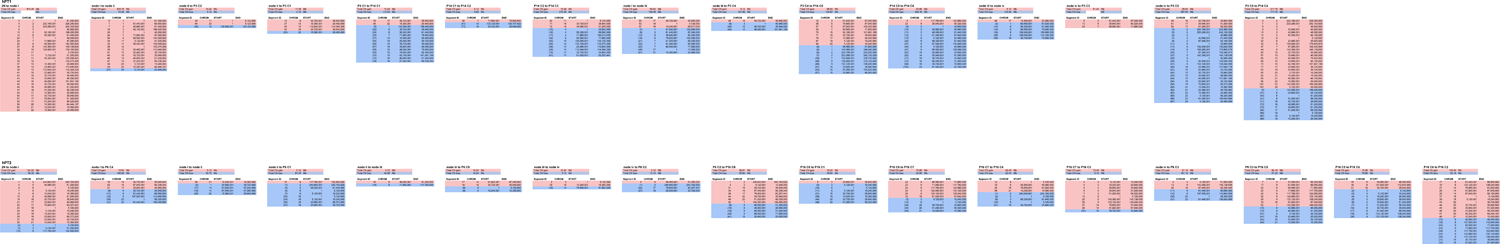
List of copy number changes that occur between the nodes of the phylogenetic trees for hPT1 and hPT2. Segment ID represents an identifier for a given segment in the genome. Copy gains are color coded in red. Copy losses are color coded in blue. Copy loss segment ID numbers are in parentheses.

### Copy number alterations in PDAC organoids translate to transcript regulation

To investigate the functionality of these copy number alterations, the transcriptomes of hPT1 and hPT2 were analyzed through RNA-seq. The gene expression fold change between the two organoids was quantified and its profile across the whole genome was averaged at every 1 Mb. CNVs between hPT1 and hPT2 were derived from the SNPa copy number profiles that also have 1 Mb resolution. The matching resolution enables the comparison between gene expression regulation and the CNVs between the two organoids (**Figure 4A**). The variation within the gene expression regulation profile is much higher than the CNV, which may be due to the trans-regulation of transcription^28^. However, the agreement between the CNV and gene expression regulation profile can be observed across the whole genome, especially where CNVs occur, such as in chromosomes 1, 3, 6, 7, 10, 14, 15, and 21. CNVs of hPT1 and hPT2 positively correlated to the gene expression regulation between the two organoids, with a statistically significant Pearson correlation coefficient of 0.42 (**Figure 4B**), suggesting “gene dosage” effects^19–21^ and functionality of chromosome copy number changes.

**Figure 4.**
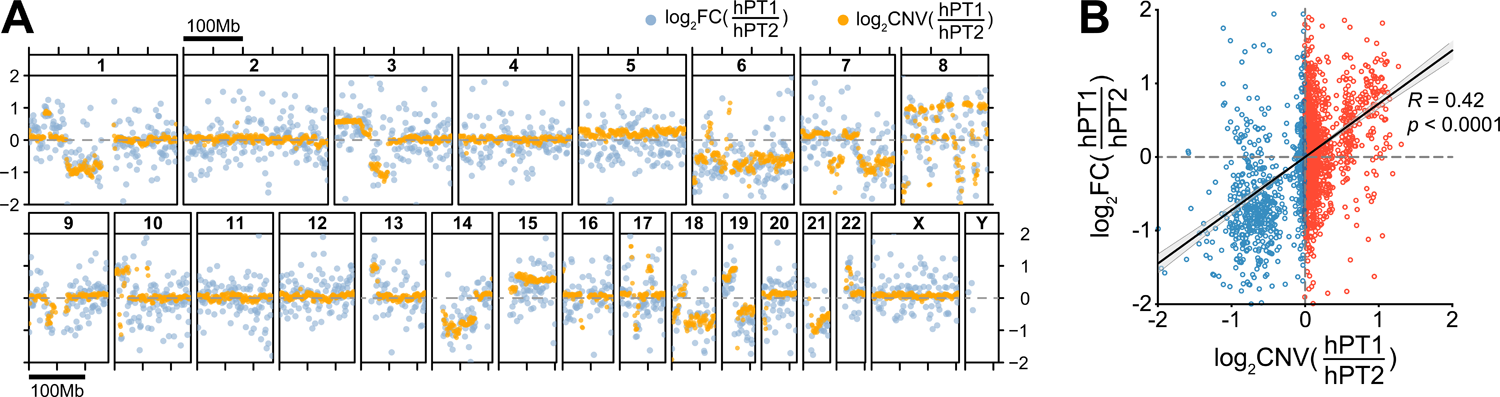
Copy number alterations translate to transcript regulations. (**A**) Both hPT1 and hPT2 organoids were subjected to RNA-seq. The gene expression fold change (log2FC, blue) between the two was averaged for every 1 Mb region across the whole genome. The plot shows good accordance between the gene expression fold change and the copy number variations (log2CNV from SNPa copy number profiles, orange) of the two organoids, especially in chromosomes 1, 3, 6, 7, 10, 14, 15, 19, and 21. (**B**) The CNV of each 1 Mb region was plotted against the corresponding gene expression fold change, resulting in a positive correlation between CNV and gene expression fold change. The correlation has a Pearson correlation coefficient (*R*) of 0.42, which is statistically significant, with *p* < 0.0001.

## Discussion

In this study, we used two independent patient-derived PDAC organoids, hPT1 and hPT2, to elucidate genomic heterogeneity within the organoid cultures. Chromosome copy number alterations were evident across the genome of both organoids (Figure 1C), and copy number gains were more prevalent than the losses, resulting in a sub-triploid ploidy. Such copy number alterations were also reported in previous PDAC genomic studies involving both tumor tissues ^9, 10^ and organoids^3, 5, 8, 18, 29^. Additionally, by performing karyotyping, one of the pioneering studies that derived the PDAC organoid^3^ also showed ploidy heterogeneity and shift within the organoid culture. Here, we further investigated this genomic heterogeneity within PDAC organoid cultures and their genomic stability with extended culture by using scWGS (Figures 2 and 3). With this technology, we can derive high-resolution copy number profiles at single-cell resolution. Clonal populations with similar copy number profiles were observed within the organoids, and the proportion of these clones was shifted with extended culture, suggesting the growth advantage of some clones. However, sub-clonal genomic heterogeneity was also observed within each clonal population, indicating the genomic instability of the PDAC cells themselves. Considering the functionality of these copy number alterations in gene expression regulation (Figure 4), we speculate that the observed genomic heterogeneity here should translate to a population of cells with heterogeneous phenotypes. Indeed, such transcriptomic heterogeneity was also reported by a single-cell RNA-seq study on PDAC organoids^30^.

All combined, the findings in this study suggest the need for future genomic studies that involve the use of PDAC organoids, to consider the genomic shift of PDAC organoids with culture. This will help to provide a more accurate interpretation of the genomic data. Furthermore, the genomic instability in PDAC organoids may not be limited to the shift in copy number profiles: It may also impact other forms of genomic variations, such as single nucleotide variations (SNVs) and other structural variants. However, the detection of SNVs from scWGS data remains challenging. This is mainly due to sequencing cost, limitations in whole genome amplification chemistry, and the availability of suitable analysis algorithms. For example, in this study, we used the multiple displacement amplification method, which introduces amplification artifacts across the genome, and the commonly used mutation detection algorithm like GATK Mutect2^31^ will detect these artifacts as SNVs (data not shown). Hence, improving genome amplification chemistry to minimize artifacts and developing algorithms that can address these amplification artifacts is needed for future scWGS studies. On another note, unlike copy number quantification, SNV analysis typically requires a read coverage of >30X, and the sequencing cost of ∼1000 cells will be very high to achieve such coverage. Alternatively, if we assume that the clones with similar copy number profiles have similar genomic variations, then we can combine their sequencing data to increase the read depth, which then enables the SNV analysis. This approach has been tested previously with some success^32^.

Future single-cell functional genomic studies are needed to elucidate the mechanistic insights behind the growth advantage of certain clones and the genomic instability within PDAC organoids. Some groups have established the technology to sequence both genomic DNA and mRNA of the same cell^33–36^, however, currently no commercially available technology enables the mass adaptation of such assays yet. Alternatively, considering the significant correlation between CNV and gene expression regulation (Figure 4), we can perform scWGS and single-cell RNA-seq independently on the same organoid line, and potentially match the copy number profile of a given clone to the corresponding gene expression profile, which then allows us to derive the key regulators of the observed genomic variations. On a different note, inference of copy number profiles from RNA-seq data has been reported in multiple studies^37–40^. However, as shown in the CNV and RNA-seq correlation in Figure 4A, the gene expression profile has much higher variations across the genome when compared to the CNV counterpart; hence, inference from RNA-seq data may lead to less accurate quantification of copy number profiles.

## Methods

### PDAC organoid culture and immunostaining

hPT1 and hPT2 PDAC organoids were derived from the primary tumors of two independent patients. The hPT1 organoids were acquired from the Barts Pancreas Tissue Bank at passage number 5 with the organoid ID B01P0735BOR (2018/14/FSU/JI/P/Organoids). The hPT2 organoids were acquired from ATCC, as part of the Human Cancer Models Initiative, at passage number 15 with the organoid name HCM-CSHL-0092-C25. The passage number listed in the manuscript represents the number of passages performed within our lab upon the first thaw. For example, hPT1 P3 in Figure 2A was passaged three times in our lab. Maintenance of PDAC organoid culture was performed following the Tuveson Laboratory Murine and Human Organoid Protocols (http://tuvesonlab.labsites.cshl.edu/wp-content/uploads/sites/49/2018/06/20170523_OrganoidProtocols.pdf), which were kindly compiled based on that lab’s their previous studies^2, 3^. For immunostaining, PDAC organoids within the Matrigel (Corning) were fixed in 4% formaldehyde (EMS) for 15 min, permeabilized by 0.5% Triton-X (Sigma) for 10 min, blocked by 5% BSA (VWR), and incubated overnight in the primary antibodies against pan-Keratin (Cell Signaling) and EpCAM (Abcam). DNA was stained with 2 µg/mL DAPI (Sigma) for 15 min. Confocal imaging was done using a Zeiss LSM 880 system with a 63X/1.4 NA oil-immersion objective. Brightfield imaging was performed using an Olympus IX71 with a digital sCMOS camera (Prime 95B, Photometrics) and a 10X/0.3 NA objective.

### Single nucleotide polymorphism microarray

DNA isolation was performed as suggested in the Tuveson Laboratory Murine and Human Organoid Protocols, except that the Blood and Cell Culture DNA Mini Kit (QIAGEN) was used instead of the suggested kit. The isolated DNA samples were sent to the Center for Applied Genomics at the Children’s Hospital of Philadelphia for the SNP array analysis using the Infinium Global Screening Array-24 v3.0 Kit (Illumina). The chromosome copy number analysis was performed as described previously^22^. Heatmaps were plotted in R by using the algorithm gtrellis v.1.28.0^41^.

### Single-cell whole-genome sequencing

For scWGS, to get a single-cell suspension, the PDAC organoids were enzymatically digested with TrypLE Express (Thermo Fisher) following the Tuveson Laboratory Murine and Human Organoid Protocols. The single-cell suspension was then processed using Chromium Single Cell CNV (10X Genomics) for scWGS. Briefly, this technology employs the droplet microfluidic method and the whole-genome multiple displacement amplification method. The resulting libraries of amplified DNA were subjected to 150-bp paired-end sequencing with a Novaseq 6000. The numbers of reads for the samples were approximately 180, 140, 420, and 530 million reads for hPT1 P3, hPT1 P14, hPT2 P6, and hPT2 P16, respectively. These read amounts provide sufficient read depth for the chromosome copy number quantification, per the manufacturer’s recommendation.

### Bioinformatics of the single-cell whole-genome sequencing data

To derive the chromosome copy number profiles, the scWGS data was analyzed using the 10x Genomics Cell Ranger DNA algorithms and visualized using the 10x Genomics Loupe scDNA Browser. The resulting copy number profiles had a resolution of 5 Mb, meaning that each chromosome was segmented into 5 Mb parts, and each segment was represented by a copy number. For the phylogenetic tree analysis, when continuous segments in a cluster had the same copy number, we merged those segments, while making sure that the merged segments could still be vertically compared to each other across all clusters. Put another way, when a cluster had different copy numbers in the segments to be merged, we shrank the number of segments until all clusters had the same copy number on the segments to be merged. We then filtered out the segments that had the same copy number across all clusters since these segments do not contribute to the heterogeneity of the cell clusters or the construction of the phylogenetic tree. In this way, we reduced the number of unnecessary segments. Next, we used PAUP^27^ to derive maximum parsimony phylogenetic trees for the early passage clusters: hPT1 P3 clusters, called T1, and hPT2 P6 clusters, called T2, respectively. Both T1 and T2 are rooted to a copy number neutral diploid genome. Under the criterion of maximum parsimony, PAUP renders a phylogenetic tree that has a minimum number of changes of copy number states for each merged segment, along with the copy number states for all the internal nodes on the tree. The distance between each cluster and the root was calculated by summing over the length of the edges along the path from the root to the cluster. To calculate the length of an edge, we first summed over the length of all segments whose copy number was different between the parent and the child nodes that the edge connects. We then divided the sum by 1 Mb. The resulting number, i.e., the copy number change measured in megabases, represents the length of this edge. We then attached the later passage clusters in hPT1 P14 to T1 through the minimal distance pairings as described in the following manner: Let set A be the clusters in hPT1 P14 that have not yet been placed on T1. Let set B be the clusters already been placed on T1. We calculated the pairwise distance between all elements in A and all elements in B. The pairwise distance was calculated in the same way as calculating the length of an edge mentioned above. Suppose pair (x, y), in which x is from A, and y is from B, has the smallest distance. We attached x to y so that x becomes a daughter node of y. Since x is attached to T1, it is no longer an element of A but becomes an element of B. We thus update sets A and B accordingly. This whole process was iterated until set A becomes empty and all clusters in hPT1 P14 are attached to T1. We repeated the whole process to attach the clusters in hPT2 P16 to T2. The resulting trees T1 and T2 are the final phylogenetic tree containing both P3 and P14 for hPT1, and P6 and P16 for hPT2. In summary, we divided the construction of the phylogenetic tree into two steps in which the first step is to construct the maximum parsimony tree from the clusters in hPT1 P3, and the clusters in hPT2 P6, whereas the second step is to attach the clusters in P14 to the tree for hPT1, and the clusters in P16 to the tree for hPT2, respectively. This way, the tree honors the different passage numbers, i.e. culturing time, of hPT1 P3 and P14, as well as hPT2 P6 and P16. The phylogenetic trees were plotted using ggtree v.3.4.0^42^ and treeio v.1.20.0^43^. Heatmap visualization of the genomic data was done using gtrellis v1.28.0^41^. The t-SNE^44^ and spectral clustering were done using the TSNE and SpectralClustering functions, respectively, in Python.

### RNA-seq sample preparation and analysis

RNA isolation was performed as suggested in the Tuveson Laboratory Murine and Human Organoid Protocols. Libraries for RNA-seq were made with the NEBNext Ultra II Directional RNA Library Prep Kit for Illumina (NEB) per the manufacturer’s instructions, followed by 150-bp paired-end sequencing with the Novaseq 6000 at Florida State University’s Translational Science Laboratory, resulting in ∼25,000,000 reads for each sample. Reads per kilobase million for each gene were calculated by normalization of the read of each gene by the sample’s total read count (in millions) and by the gene length (in kilobases). The gene expression profile was plotted in R by using the algorithm gtrellis v.1.28.0^41^.

## Acknowledgment

The authors would like to thank David A. Tuveson (Cold Spring Harbor Laboratory) and Herve Tiriac (University of California San Diego) for their support and guidance in PDAC organoid culture. The authors would like to thank Dr. Terra Bradley for the careful editing of the manuscript. The authors in this study were supported by startup funds from Florida State University, an award from the Florida Department of Health’s Bankhead-Coley Cancer Research Program (award number 21B11), and a collaborative Trans-Network Project linked to the Physical Science-Oncology Project 5U01 CA214282 from the National Cancer Institute of the National Institutes of Health. The hPT1 PDAC organoid line is from the Barts Pancreas Tissue Bank, a research tissue bank at the Barts Cancer Institute, Queen Mary University of London, London, UK, https://www.bartspancreastissuebank.org.uk/index.html, and funded by Pancreatic Cancer Research Fund. Ahmet Imrali was instrumental in developing the organoids at the Barts Pancreas Tissue Bank. The hPT2 PDAC organoid line is from the Human Cancer Models Initiative, https://ocg.cancer.gov/programs/HCMI.

## Notes

### Competing Interest Statement

The authors have declared no competing interest.

### Summary of Updates

Funding sources in acknowledgment updated; Manuscript is double spaced; Figures rearranged to the back of the manuscript; ORCID linked to authors.

## Reference

1. American Cancer Society. Cancer Facts & Figures 2022. Atlanta: American Cancer Society, 2022.

2. Huch M, Bonfanti P, Boj SF, Sato T, Loomans CJ, van de Wetering M, Sojoodi M, Li VS, Schuijers J, Gracanin A, Ringnalda F, Begthel H, Hamer K, Mulder J, van Es JH, de Koning E, Vries RG, Heimberg H & Clevers H. Unlimited in vitro expansion of adult bi-potent pancreas progenitors through the Lgr5/R-spondin axis. EMBO J 2013; 32: 2708–21.

3. Boj SF, Hwang CI, Baker LA, Chio, II, Engle DD, Corbo V, Jager M, Ponz-Sarvise M, Tiriac H, Spector MS, Gracanin A, Oni T, Yu KH, van Boxtel R, Huch M, Rivera KD, Wilson JP, Feigin ME, Ohlund D, Handly-Santana A, Ardito-Abraham CM, Ludwig M, Elyada E, Alagesan B, Biffi G, Yordanov GN, Delcuze B, Creighton B, Wright K, Park Y, Morsink FH, Molenaar IQ, Borel Rinkes IH, Cuppen E, Hao Y, Jin Y, Nijman IJ, Iacobuzio-Donahue C, Leach SD, Pappin DJ, Hammell M, Klimstra DS, Basturk O, Hruban RH, Offerhaus GJ, Vries RG, Clevers H & Tuveson DA. Organoid models of human and mouse ductal pancreatic cancer. Cell 2015; 160: 324–38.

4. Tiriac H, Belleau P, Engle DD, Plenker D, Deschenes A, Somerville TDD, Froeling FEM, Burkhart RA, Denroche RE, Jang GH, Miyabayashi K, Young CM, Patel H, Ma M, LaComb JF, Palmaira RLD, Javed AA, Huynh JC, Johnson M, Arora K, Robine N, Shah M, Sanghvi R, Goetz AB, Lowder CY, Martello L, Driehuis E, LeComte N, Askan G, Iacobuzio-Donahue CA, Clevers H, Wood LD, Hruban RH, Thompson E, Aguirre AJ, Wolpin BM, Sasson A, Kim J, Wu M, Bucobo JC, Allen P, Sejpal DV, Nealon W, Sullivan JD, Winter JM, Gimotty PA, Grem JL, DiMaio DJ, Buscaglia JM, Grandgenett PM, Brody JR, Hollingsworth MA, O’Kane GM, Notta F, Kim E, Crawford JM, Devoe C, Ocean A, Wolfgang CL, Yu KH, Li E, Vakoc CR, Hubert B, Fischer SE, Wilson JM, Moffitt R, Knox J, Krasnitz A, Gallinger S & Tuveson DA. Organoid Profiling Identifies Common Responders to Chemotherapy in Pancreatic Cancer. Cancer Discov 2018; 8: 1112–1129.

5. Driehuis E, van Hoeck A, Moore K, Kolders S, Francies HE, Gulersonmez MC, Stigter ECA, Burgering B, Geurts V, Gracanin A, Bounova G, Morsink FH, Vries R, Boj S, van Es J, Offerhaus GJA, Kranenburg O, Garnett MJ, Wessels L, Cuppen E, Brosens LAA & Clevers H. Pancreatic cancer organoids recapitulate disease and allow personalized drug screening. Proc Natl Acad Sci U S A 2019;

6. Huang L, Holtzinger A, Jagan I, BeGora M, Lohse I, Ngai N, Nostro C, Wang R, Muthuswamy LB, Crawford HC, Arrowsmith C, Kalloger SE, Renouf DJ, Connor AA, Cleary S, Schaeffer DF, Roehrl M, Tsao MS, Gallinger S, Keller G & Muthuswamy SK. Ductal pancreatic cancer modeling and drug screening using human pluripotent stem cell- and patient-derived tumor organoids. Nat Med 2015; 21: 1364–71.

7. Walsh AJ, Castellanos JA, Nagathihalli NS, Merchant NB & Skala MC. Optical Imaging of Drug-Induced Metabolism Changes in Murine and Human Pancreatic Cancer Organoids Reveals Heterogeneous Drug Response. Pancreas 2016; 45: 863–9.

8. Seino T, Kawasaki S, Shimokawa M, Tamagawa H, Toshimitsu K, Fujii M, Ohta Y, Matano M, Nanki K, Kawasaki K, Takahashi S, Sugimoto S, Iwasaki E, Takagi J, Itoi T, Kitago M, Kitagawa Y, Kanai T & Sato T. Human Pancreatic Tumor Organoids Reveal Loss of Stem Cell Niche Factor Dependence during Disease Progression. Cell Stem Cell 2018; 22: 454–467 e6.

9. Campbell PJ, Yachida S, Mudie LJ, Stephens PJ, Pleasance ED, Stebbings LA, Morsberger LA, Latimer C, McLaren S, Lin ML, McBride DJ, Varela I, Nik-Zainal SA, Leroy C, Jia M, Menzies A, Butler AP, Teague JW, Griffin CA, Burton J, Swerdlow H, Quail MA, Stratton MR, Iacobuzio-Donahue C & Futreal PA. The patterns and dynamics of genomic instability in metastatic pancreatic cancer. Nature 2010; 467: 1109–13.

10. Makohon-Moore AP, Zhang M, Reiter JG, Bozic I, Allen B, Kundu D, Chatterjee K, Wong F, Jiao Y, Kohutek ZA, Hong J, Attiyeh M, Javier B, Wood LD, Hruban RH, Nowak MA, Papadopoulos N, Kinzler KW, Vogelstein B & Iacobuzio-Donahue CA. Limited heterogeneity of known driver gene mutations among the metastases of individual patients with pancreatic cancer. Nat Genet 2017; 49: 358–366.

11. Yachida S, Jones S, Bozic I, Antal T, Leary R, Fu B, Kamiyama M, Hruban RH, Eshleman JR, Nowak MA, Velculescu VE, Kinzler KW, Vogelstein B & Iacobuzio-Donahue CA. Distant metastasis occurs late during the genetic evolution of pancreatic cancer. Nature 2010; 467: 1114–7.

12. Hayashi A, Ho Y-j, Makohon-Moore AP, Zucker A, Hong J, Reiter JG, Huang J, Zhang L, Attiyeh MA, Baez P, Kappagantula R, Melchor JP, O’Reilly EM, Socci ND, Oki S, Lowe SW & Iacobuzio-Donahue CA. Evolutionary Dynamics of Non-Coding Regions in Pancreatic Ductal Adenocarcinoma. bioRxiv 2020; 2020.09.11.294389.

13. Peng J, Sun BF, Chen CY, Zhou JY, Chen YS, Chen H, Liu L, Huang D, Jiang J, Cui GS, Yang Y, Wang W, Guo D, Dai M, Guo J, Zhang T, Liao Q, Liu Y, Zhao YL, Han DL, Zhao Y, Yang YG & Wu W. Single-cell RNA-seq highlights intra-tumoral heterogeneity and malignant progression in pancreatic ductal adenocarcinoma. Cell Res 2019; 29: 725–738.

14. Maddipati R & Stanger BZ. Pancreatic Cancer Metastases Harbor Evidence of Polyclonality. Cancer Discov 2015; 5: 1086–97.

15. Harada T, Okita K, Shiraishi K, Kusano N, Kondoh S & Sasaki K. Interglandular cytogenetic heterogeneity detected by comparative genomic hybridization in pancreatic cancer. Cancer Res 2002; 62: 835–9.

16. Le Large TY, Mantini G, Meijer LL, Pham TV, Funel N, van Grieken NC, Kok B, Knol J, van Laarhoven HW, Piersma SR, Jimenez CR, Kazemier G, Giovannetti E & Bijlsma MF. Microdissected pancreatic cancer proteomes reveal tumor heterogeneity and therapeutic targets. JCI Insight 2020; 5:

17. Cancer Genome Atlas Research Network. Electronic address aadhe & Cancer Genome Atlas Research N. Integrated Genomic Characterization of Pancreatic Ductal Adenocarcinoma. Cancer Cell 2017; 32: 185–203 e13.

18. Miyabayashi K, Baker LA, Deschenes A, Traub B, Caligiuri G, Plenker D, Alagesan B, Belleau P, Li S, Kendall J, Jang GH, Kawaguchi RK, Somerville TDD, Tiriac H, Hwang CI, Burkhart RA, Roberts NJ, Wood LD, Hruban RH, Gillis J, Krasnitz A, Vakoc CR, Wigler M, Notta F, Gallinger S, Park Y & Tuveson DA. Intraductal Transplantation Models of Human Pancreatic Ductal Adenocarcinoma Reveal Progressive Transition of Molecular Subtypes. Cancer Discov 2020; 10: 1566–1589.

19. Clemente-Ruiz M, Murillo-Maldonado JM, Benhra N, Barrio L, Perez L, Quiroga G, Nebreda AR & Milan M. Gene Dosage Imbalance Contributes to Chromosomal Instability-Induced Tumorigenesis. Dev Cell 2016; 36: 290–302.

20. Medina-Martinez I, Barron V, Roman-Bassaure E, Juarez-Torres E, Guardado-Estrada M, Espinosa AM, Bermudez M, Fernandez F, Venegas-Vega C, Orozco L, Zenteno E, Kofman S & Berumen J. Impact of gene dosage on gene expression, biological processes and survival in cervical cancer: a genome-wide follow-up study. PLoS One 2014; 9: e97842.

21. Rice AM & McLysaght A. Dosage-sensitive genes in evolution and disease. BMC Biol 2017; 15: 78.

22. Irianto J, Xia Y, Pfeifer CR, Athirasala A, Ji J, Alvey C, Tewari M, Bennett RR, Harding SM, Liu AJ, Greenberg RA & Discher DE. DNA Damage Follows Repair Factor Depletion and Portends Genome Variation in Cancer Cells after Pore Migration. Curr Biol 2017; 27: 210–223.

23. Carter NP. Methods and strategies for analyzing copy number variation using DNA microarrays. Nat Genet 2007; 39: S16–21.

24. Stephens PJ, Greenman CD, Fu B, Yang F, Bignell GR, Mudie LJ, Pleasance ED, Lau KW, Beare D, Stebbings LA, McLaren S, Lin ML, McBride DJ, Varela I, Nik-Zainal S, Leroy C, Jia M, Menzies A, Butler AP, Teague JW, Quail MA, Burton J, Swerdlow H, Carter NP, Morsberger LA, Iacobuzio-Donahue C, Follows GA, Green AR, Flanagan AM, Stratton MR, Futreal PA & Campbell PJ. Massive genomic rearrangement acquired in a single catastrophic event during cancer development. Cell 2011; 144: 27–40.

25. Waddell N, Pajic M, Patch AM, Chang DK, Kassahn KS, Bailey P, Johns AL, Miller D, Nones K, Quek K, Quinn MC, Robertson AJ, Fadlullah MZ, Bruxner TJ, Christ AN, Harliwong I, Idrisoglu S, Manning S, Nourse C, Nourbakhsh E, Wani S, Wilson PJ, Markham E, Cloonan N, Anderson MJ, Fink JL, Holmes O, Kazakoff SH, Leonard C, Newell F, Poudel B, Song S, Taylor D, Waddell N, Wood S, Xu Q, Wu J, Pinese M, Cowley MJ, Lee HC, Jones MD, Nagrial AM, Humphris J, Chantrill LA, Chin V, Steinmann AM, Mawson A, Humphrey ES, Colvin EK, Chou A, Scarlett CJ, Pinho AV, Giry-Laterriere M, Rooman I, Samra JS, Kench JG, Pettitt JA, Merrett ND, Toon C, Epari K, Nguyen NQ, Barbour A, Zeps N, Jamieson NB, Graham JS, Niclou SP, Bjerkvig R, Grutzmann R, Aust D, Hruban RH, Maitra A, Iacobuzio-Donahue CA, Wolfgang CL, Morgan RA, Lawlor RT, Corbo V, Bassi C, Falconi M, Zamboni G, Tortora G, Tempero MA, Australian Pancreatic Cancer Genome I, Gill AJ, Eshleman JR, Pilarsky C, Scarpa A, Musgrove EA, Pearson JV, Biankin AV & Grimmond SM. Whole genomes redefine the mutational landscape of pancreatic cancer. Nature 2015; 518: 495–501.

26. Notta F, Chan-Seng-Yue M, Lemire M, Li Y, Wilson GW, Connor AA, Denroche RE, Liang SB, Brown AM, Kim JC, Wang T, Simpson JT, Beck T, Borgida A, Buchner N, Chadwick D, Hafezi-Bakhtiari S, Dick JE, Heisler L, Hollingsworth MA, Ibrahimov E, Jang GH, Johns J, Jorgensen LG, Law C, Ludkovski O, Lungu I, Ng K, Pasternack D, Petersen GM, Shlush LI, Timms L, Tsao MS, Wilson JM, Yung CK, Zogopoulos G, Bartlett JM, Alexandrov LB, Real FX, Cleary SP, Roehrl MH, McPherson JD, Stein LD, Hudson TJ, Campbell PJ & Gallinger S. A renewed model of pancreatic cancer evolution based on genomic rearrangement patterns. Nature 2016; 538: 378–382.

27. Swofford DL. PAUP*. Phylogenetic Analysis Using Parsimony (*and Other Methods). Version 4.0a. Sinauer Associates, Sunderland, Massachusetts (2003).

28. Jaenisch R & Bird A. Epigenetic regulation of gene expression: how the genome integrates intrinsic and environmental signals. Nat Genet 2003; 33 Suppl: 245-54.

29. Gendoo DMA, Denroche RE, Zhang A, Radulovich N, Jang GH, Lemire M, Fischer S, Chadwick D, Lungu IM, Ibrahimov E, Cao PJ, Stein LD, Wilson JM, Bartlett JMS, Tsao MS, Dhani N, Hedley D, Gallinger S & Haibe-Kains B. Whole genomes define concordance of matched primary, xenograft, and organoid models of pancreas cancer. PLoS Comput Biol 2019; 15: e1006596.

30. Krieger TG, Le Blanc S, Jabs J, Ten FW, Ishaque N, Jechow K, Debnath O, Leonhardt CS, Giri A, Eils R, Strobel O & Conrad C. Single-cell analysis of patient-derived PDAC organoids reveals cell state heterogeneity and a conserved developmental hierarchy. Nat Commun 2021; 12: 5826.

31. Auwera GAVd & O’Connor BD. Genomics in the Cloud: Using Docker, GATK, and WDL in Terra. O’Reilly Media, 2020.

32. Velazquez-Villarreal EI, Maheshwari S, Sorenson J, Fiddes IT, Kumar V, Yin Y, Webb MG, Catalanotti C, Grigorova M, Edwards PA, Carpten JD & Craig DW. Single-cell sequencing of genomic DNA resolves sub-clonal heterogeneity in a melanoma cell line. Commun Biol 2020; 3: 318.

33. Zachariadis V, Cheng H, Andrews N & Enge M. A Highly Scalable Method for Joint Whole-Genome Sequencing and Gene-Expression Profiling of Single Cells. Mol Cell 2020; 80: 541–553 e5.

34. Dey SS, Kester L, Spanjaard B, Bienko M & van Oudenaarden A. Integrated genome and transcriptome sequencing of the same cell. Nat Biotechnol 2015; 33: 285–289.

35. Macaulay IC, Haerty W, Kumar P, Li YI, Hu TX, Teng MJ, Goolam M, Saurat N, Coupland P, Shirley LM, Smith M, Van der Aa N, Banerjee R, Ellis PD, Quail MA, Swerdlow HP, Zernicka-Goetz M, Livesey FJ, Ponting CP & Voet T. G&T-seq: parallel sequencing of single-cell genomes and transcriptomes. Nat Methods 2015; 12: 519–22.

36. Hou Y, Guo H, Cao C, Li X, Hu B, Zhu P, Wu X, Wen L, Tang F, Huang Y & Peng J. Single-cell triple omics sequencing reveals genetic, epigenetic, and transcriptomic heterogeneity in hepatocellular carcinomas. Cell Res 2016; 26: 304–19.

37. Gao R, Bai S, Henderson YC, Lin Y, Schalck A, Yan Y, Kumar T, Hu M, Sei E, Davis A, Wang F, Shaitelman SF, Wang JR, Chen K, Moulder S, Lai SY & Navin NE. Delineating copy number and clonal substructure in human tumors from single-cell transcriptomes. Nat Biotechnol 2021; 39: 599–608.

38. Serin Harmanci A, Harmanci AO & Zhou X. CaSpER identifies and visualizes CNV events by integrative analysis of single-cell or bulk RNA-sequencing data. Nat Commun 2020; 11: 89.

39. Fan J, Lee HO, Lee S, Ryu DE, Lee S, Xue C, Kim SJ, Kim K, Barkas N, Park PJ, Park WY & Kharchenko PV. Linking transcriptional and genetic tumor heterogeneity through allele analysis of single-cell RNA-seq data. Genome Res 2018; 28: 1217–1227.

40. Tickle T, Tirosh I, Georgescu C, Brown M & Haas B. inferCNV of the Trinity CTAT Project. Klarman Cell Observatory, Broad Institute of MIT and Harvard, Cambridge, MA, USA (2019).https://github.com/broadinstitute/inferCNV

41. Gu Z, Eils R & Schlesner M. gtrellis: an R/Bioconductor package for making genome-level Trellis graphics. BMC Bioinformatics 2016; 17: 169.

42. Yu G. Using ggtree to Visualize Data on Tree-Like Structures. Curr Protoc Bioinformatics 2020; 69: e96.

43. Wang LG, Lam TT, Xu S, Dai Z, Zhou L, Feng T, Guo P, Dunn CW, Jones BR, Bradley T, Zhu H, Guan Y, Jiang Y & Yu G. Treeio: An R Package for Phylogenetic Tree Input and Output with Richly Annotated and Associated Data. Mol Biol Evol 2020; 37: 599–603.

44. Maaten Lvd & Hinton G. Visualizing Data using t-SNE. Journal of Machine Learning Research 2008; 9: 2579–2605.

